# Extended-Spectrum and Last-Resort β-Lactams Exert Increasing Impacts on Chicken Manure-Derived Copiotrophic Microbiomes and Resistomes

**DOI:** 10.64898/2026.02.01.700757

**Authors:** Chagai Davidovich, Shlomo E Blum, Eddie Cytryn

## Abstract

Understanding the environmental dissemination of antimicrobial resistance (AMR) is crucial for mitigating its spread and for reducing the clinical burden of resistant infections. Within a One Health framework, animal husbandry systems, and particularly animal gut microbiomes, are widely used as models for agricultural environments, because their high densities and recurrent antibiotic exposure create strong selective pressures that can impact both environmental and human resistomes. In a previous study, we demonstrated that copiotrophic enrichment in medium that stimulates growth of gut-associated bacteria, can reveal clinically associated bacterial populations and associated antibiotic resistance genes (ARGs) that are not detected by direct environmental sampling. In the present study, we applied this enrichment approach to poultry manure to examine how β-lactamase-resistant antibiotics (cefotaxime and meropenem) shape copiotrophic gut-like microbial community composition and resistance profiles relative to an earlier β-lactam (ampicillin) and to antibiotic-free controls. Our results show that β-lactamase-resistant antibiotics induced more pronounced shifts in both microbial diversity and ARG profiles than the early β-lactam. When focusing on specific taxa, differential responses were observed correlating to antimicrobial range of action. Together, these findings underline how the diversity of pathogen and AMR indicators in gut microbiomes may be shaped by different selective pressures imposed by β-lactamase-resistant antibiotics.

## Introduction

Antimicrobial resistance (AMR) has become a major medical threat due to treatment failure caused by the continued rise of multidrug-resistant infections (Naghavi et al., 2024), and is therefore increasingly studied within the One Health framework (Berendonk et al., 2015. Agroecosystems and agronomic practices such as extensive use of antibiotics in food-animal production and amendment of animal manure for soil fertilization, create AMR hotspots, leading to dissemination of antibiotic-resistant bacteria (ARB) and their antibiotic resistance genes (ARGs) (Larsson & Flach, 2022a).

While the spread of AMR in clinical settings has been extensively studied, the environmental dimensions, and specifically the contribution of animal husbandry to the dissemination of AMR determinants, remain less understood (Iwu et al., 2020). Food production animals carry a variety of AMR determinants which can be transmitted to humans via direct consumption or when animal manure is used for fertilization (Heuer et al., 2011; Muurinen et al., 2017; Zalewska et al., 2021). The prophylactic and therapeutic use of antibiotics in animal husbandry, notably in poultry farming, can promote the maintenance and dissemination of resistance determinants through co-selection mechanisms, even when the corresponding antibiotic classes are not directly used (Gupta et al., 2021; Larsson & Flach, 2022b).

β-lactams are a diverse class of antibiotics that are widely used in human and veterinary medicine, playing a central role in treating infections caused by clinically relevant Gram-positive and Gram-negative bacterial pathogens (Bush, 2018). Most β-lactam resistance is caused by β-lactamases, which are frequently encoded on conjugative plasmids that are horizontally transferred between bacteria, facilitating the emergence of resistance in pathogens (Bush & Bradford, 2020). Resistance to early β-lactam antibiotics (e.g., ampicillin and penicillin) due to first-generation β-lactamases prompted for the development and introduction of novel β-lactamase-resistant compounds including resistance to extended-spectrum β-lactamases (second and third-generation cephalosporins), and subsequently of last-resort β-lactams (carbapenems) (Essack, 2001). Increasing bacterial resistance also to these new antibiotics underscores the need for novel therapeutic strategies (Codjoe & Donkor, 2017), but no less importantly, the need for continuous monitoring and identification of AMR dissemination pathways and detection of novel AMR variants.

In recent years, there is a growing realization that tackling AMR in the environment necessitates high sensitivity detection methods, because even small loads of ARB and associated ARGs can proliferate following exposure to favorable conditions, especially under selective pressure (Fortunato et al., 2018). We recently developed a copiotrophic enrichment platform that facilitates the detection of clinically relevant ARB and ARGs which persist below limits of detection (LOD) in anthropogenically contaminated soil and produce (Marano et al., 2021). Previous reports showed that cultivation in Brain Heart Infusion (BHI) strongly promotes the growth of commensal gut-associated bacteria (Yousi et al., 2019a). Applying BHI to our enrichment platform enabled detection of clinically relevant ARB and ARGs including extended-spectrum β-lactamase (ESBL)-producing *E. coli*, which were below LOD in direct analyses, as well as other pathogen indicators, commensals and AMR determinants endemic to gut microbiomes (Davidovich et al., 2025).

The objective of this study was to evaluate how early β-lactams and β-lactamase-resistant antibiotics affect the taxonomic structure and ARG composition of gut-like bacterial communities. A simplified reconstructed gut-like microbial community was established through BHI-based copiotrophic enrichment using poultry manure as the inoculum, and a combination of cultivation-based approaches, amplicon sequencing, long-read shotgun metagenomics, and advanced bioinformatic analyses was employed to characterize bacterial community structure and ARG composition, providing comprehensive insight into gut microbiome and resistome dynamics following antibiotic exposure in the gut.

## Results

### Ampicillin had a minor impact on bacterial community composition, whereas cefotaxime and meropenem caused pronounced shifts

Manure samples from laying hens were collected and enriched overnight in anaerobic BHI medium without antibiotics (enrichment phase 1, EP-1). These enrichments were then used as inocula for a second BHI enrichment (enrichment phase 2, EP-2) without antibiotics (no-AB), or BHI supplemented with one of three β-lactam antibiotics: ampicillin (8 mg/L), cefotaxime (2 mg/L), or meropenem (8 mg/L). Antibiotic concentrations were selected based on minimum inhibitory concentration (MIC) clinical breakpoints as defined by the European Committee on Antimicrobial Susceptibility Testing (EUCAST) for Enterobacterales (EUCAST, 2021).

EP-1 and EP-2 showed reduced bacterial diversity, and a marked shift in composition relative to the native manure samples (pre-enrichment) (p < 0.01), however, in EP-2, β-lactam supplements did not lower the diversity further in comparison to the antibiotic-free control (Fig. 1A).

**Fig 1:**
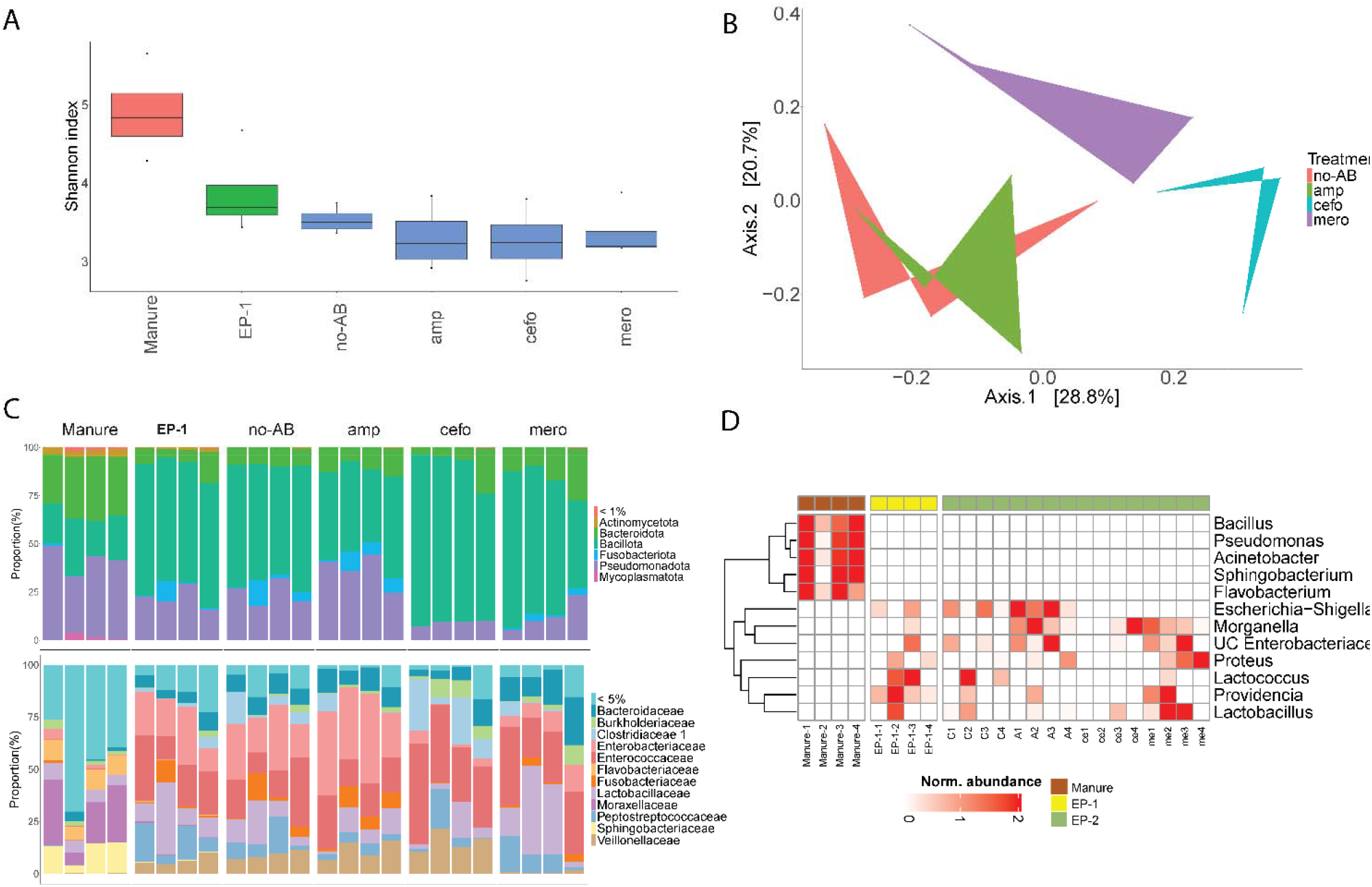
Impact of β-lactam exposure on bacterial communities based on 16S rRNA amplicon sequencing. A) α-diversity estimated based on the Shannon diversity index (linear mixed-effects models p.value < 0.05). Blue bars represent EP-2 samples. B) PCoA of β-diversity based on Bray-Curtis dissimilarity distances of EP-2 enrichments (linear mixed-effects models p.value < 0.05). C) Community composition at the phylum (upper) and family (lower) levels of original manure, EP-1, and EP-2 (represented by the applied antibiotics). D) Relative abundance of genera that are differentially abundant between manure and EP-1 and EP-2 samples (MaAsLin2 * qvalue < 0.05 |coefficient| > 1). EP-1: Enrichment phase 1, EP-2: Enrichment phase 2. No-AB: antibiotic-free control; amp: ampicillin; cefo: cefotaxime; mero: meropenem.

Generally, enrichments (both EP-1 and EP-2) led to proliferation of enteric genera such as *Providencia, Morganella*, and unclassified *Enterobacteriaceae* (Fig 1D). Surprisingly, the microbiome following ampicillin treatment was not significantly different from the antibiotic-free control. In contrast, cefotaxime and meropenem supplementation resulted in significant shifts in microbiome composition, with reduced relative abundance of Pseudomonadota and increased abundance of Bacillota. At the family level, these treatments were also associated with a decreased representation of *Enterobacteriaceae* (p < 0.05) (Fig. 1B, C, supplementary Fig 1).

### Bacterial responses to β-lactam exposure

Differential abundance analysis of the EP-2 16S rRNA gene amplicon data revealed significant differences in the occurrence of genera containing clinically relevant strains. Specifically, *Escherichia-Shigella* were more abundant in no-AB and ampicillin treatments compared to cefotaxime and meropenem treatments. The relative abundance of *Lactococcus* and Clostridia-associated genera were significantly lower in the ampicillin treatment compared to the no-AB enrichment, despite the overall similarity in bacterial composition between no-AB and the ampicillin enrichment. In general, the relative abundances of most pathogen-associated genera were negatively affected by all three β-lactams, and cefotaxime had the most significant impact. Nonetheless, the relative abundance of *Sutterella, Lachnoclostridium*, and *Enterococcus* increased in the cefotaxime treatments, while the relative abundance of *Providencia* was higher in the meropenem treatment and lower in the cefotaxime-treated samples, and that of *Proteus* decreased only in the cefotaxime treatment (MaAsLin2: -1 < coefficient < 1, qvalue < 0.05) (Fig. 2 and supplementary Fig. 2).

**Fig 2:**
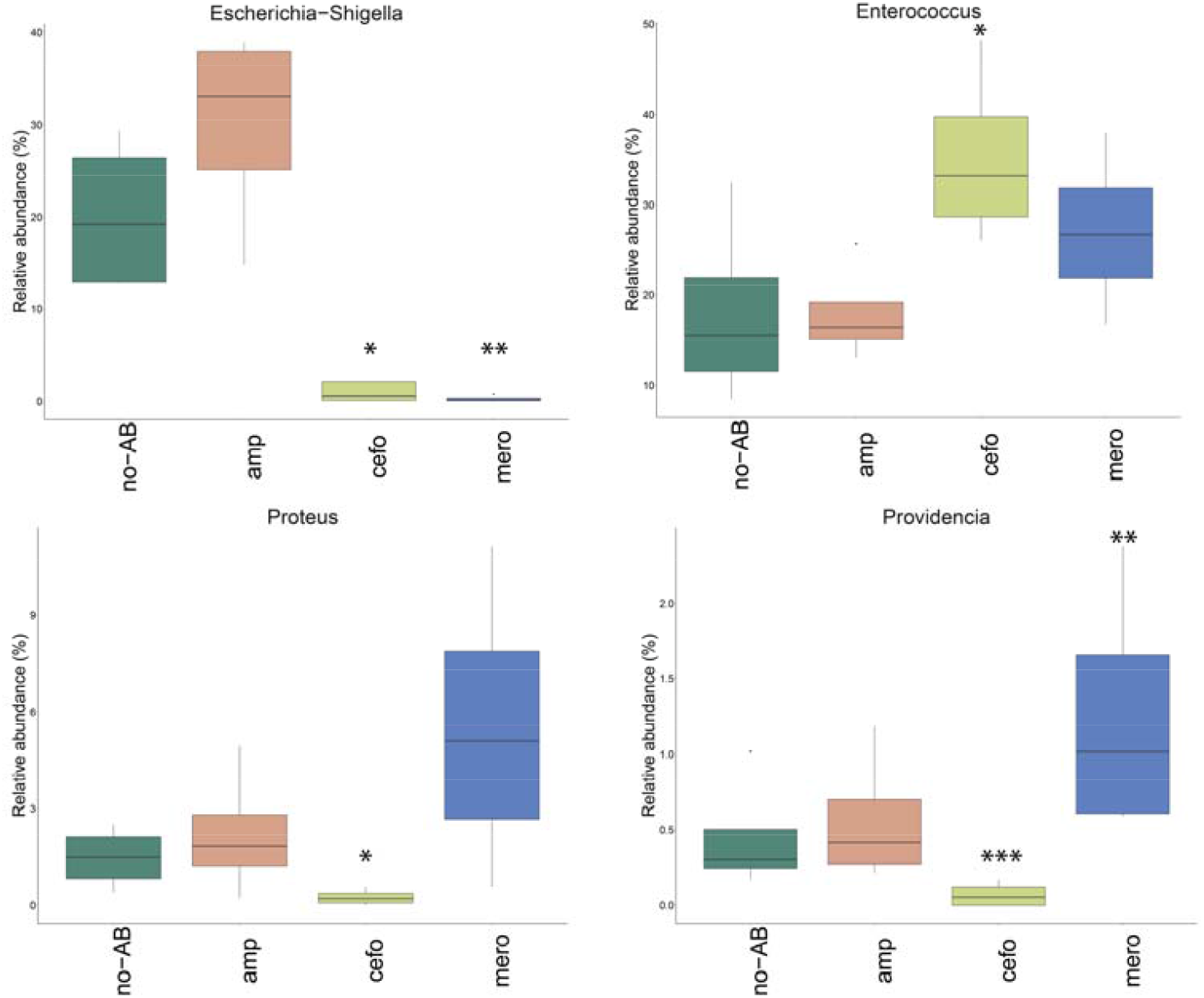
Relative abundance of genera that are differentially abundant in EP-2 enrichments without antibiotics and in enrichments supplemented with different β-lactams (MaAsLin2 * qvalue < 0.05, ** qvalue < 0.005, *** qvalue < 0.0005, |coefficient| > 1). No-AB: antibiotic-free control; amp: ampicillin; cefo: cefotaxime; mero: meropenem.

Using selective medium, approximately 10□ CFU/mL coliforms were quantified in both EP-1 and the no-AB EP-2 samples. These levels were slightly lower in the EP-2 ampicillin treatments, but markedly decreased in the cefotaxime treatments, in which no growth was observed in two of the four replicates, and reduced growth (∼10□ CFU/mL) occurred in the remaining two. Coliforms did not grow in any of the meropenem-supplemented enrichments (Dunnett□s test p.value < 0.05) (Fig. 3). MALDI-TOF analysis for coliform identification revealed that all isolates belonged to the *Escherichia–Shigella* genus, consistent with the prevalence of *Escherichia–Shigella* observed in the microbiome analyses described above, underlining the fact that in the tested poultry manure microbiome a large fraction of these pathogen indicators are resistant to ampicillin, a smaller proportion are resistant to cefotaxime and nearly all are sensitive to meropenem.

**Fig 3:**
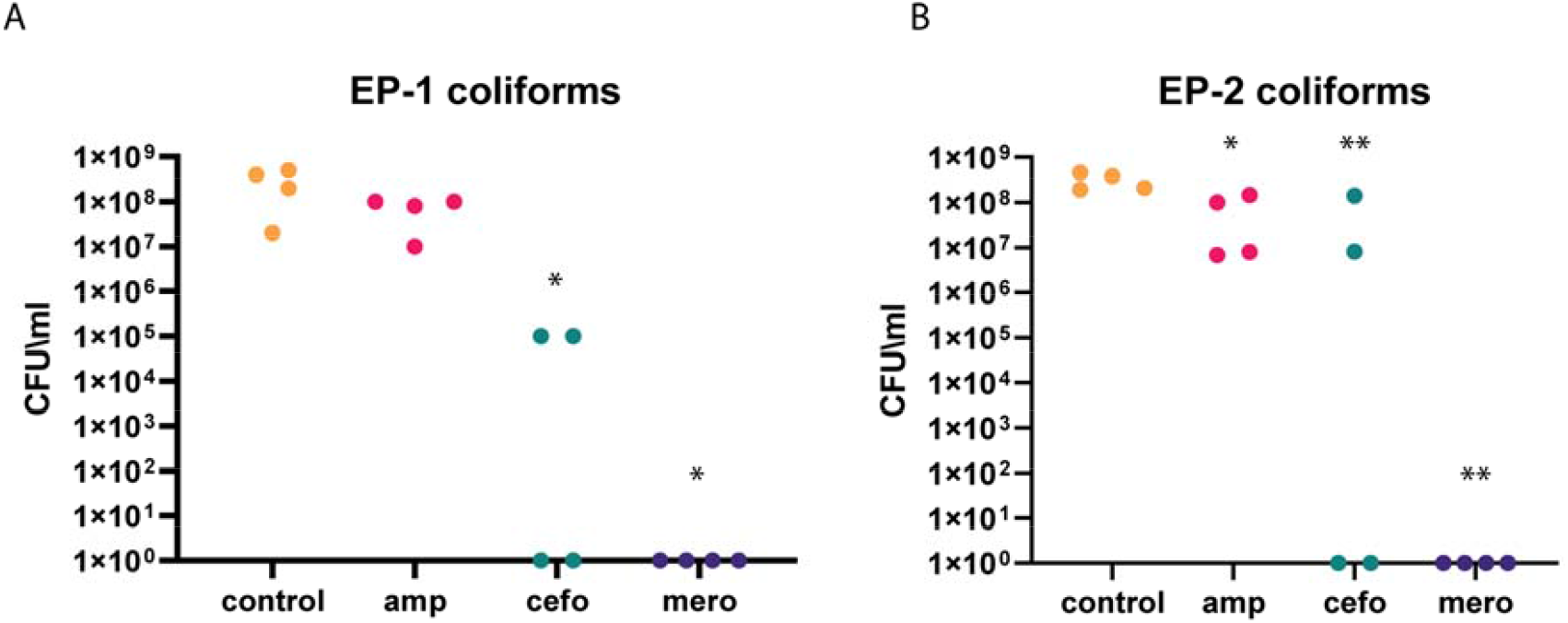
Quantification of fecal coliforms in EP-1 (A) and EP-2 (B) samples. EP-1 samples were plated on CCA, whereas the EP2 enrichment samples were plated on CCA supplemented with the indicated antibiotics (Dunnet’s test * p.value < 0.05 ** p.value < 0.005).

### Resistomes and mobilomes are significantly modulated following exposure to cefotaxime and meropenem, but not by ampicillin

ARGs and MGEs were extrapolated from shotgun metagenomic data to evaluate the impact of β-lactam application on AMR determinants. No significant differences were observed in α-diversity between the EP-1 and the no-AB and ampicillin-supplemented EP-2 samples. In contrast, cefotaxime- and meropenem-supplemented EP-2 showed a marked reduction in diversity of both ARGs and MGEs (Shannon index, p < 0.05) (Fig 4A, B). This included a significant decrease in the relative abundance of the *E. coli*-associated *ampC* gene, and in most multidrug resistance (MDR) efflux pumps. A similar trend was observed for the plasmid mediated quinolone resistance gene *qnrB72*, although the relative abundance of *qnrD2* increased in meropenem-treated samples. The relative abundance of the macrolide resistance gene *lsaA* was lower in the EP-2 ampicillin samples than the no-AB samples. In cefotaxime-supplemented samples, the macrolide efflux gene *macB* decreased, whereas the aminoglycoside resistance gene *AAC(6*^′^*)-Iih* increased. Additionally, the relative abundance of the polymyxin resistance gene *rosB* and the tetracycline resistance gene *tetA* were both lower in the meropenem enrichments than in the other EP-2 samples (Fig. 4C).

**Fig 4:**
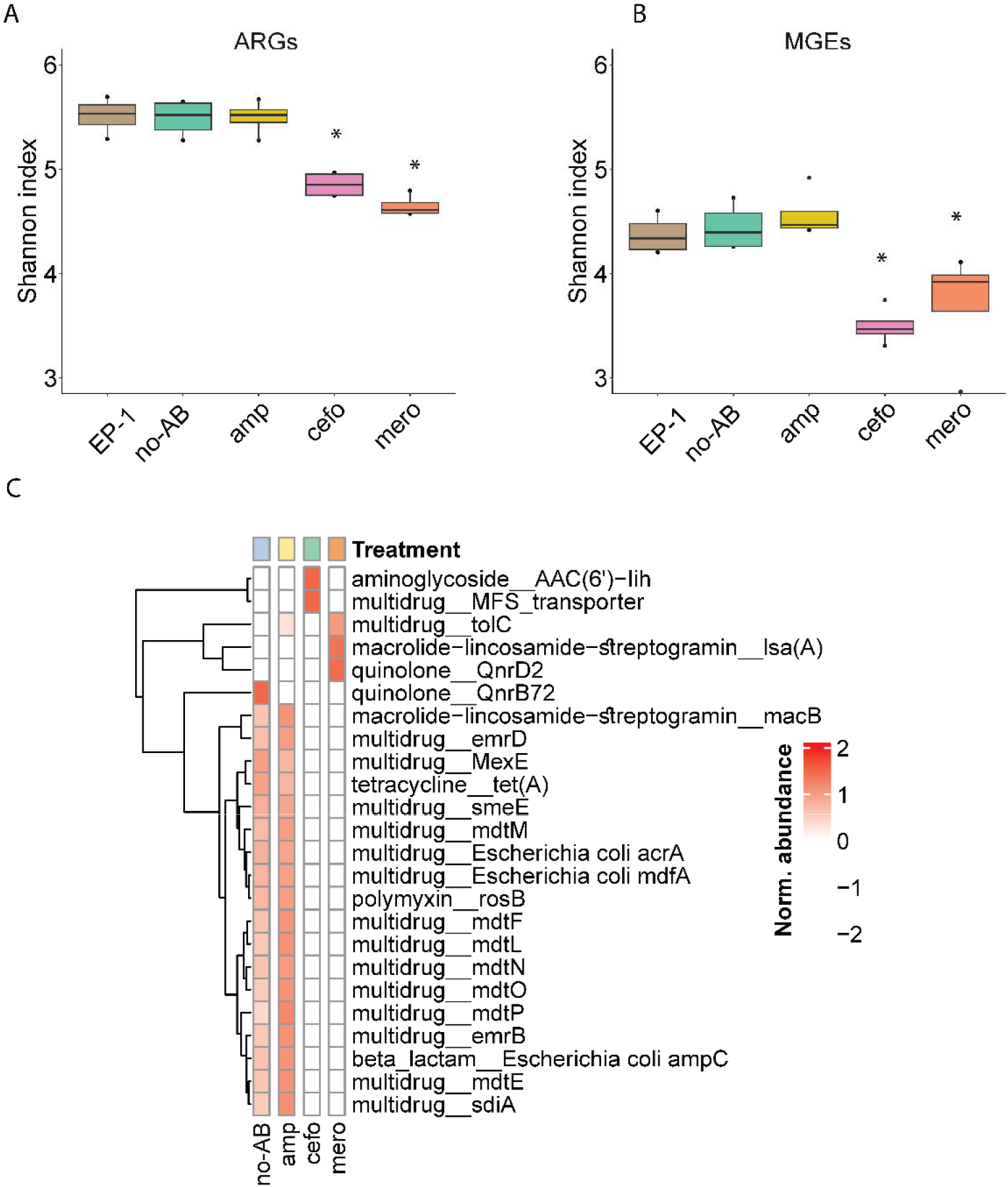
ARG diversity and composition across treatments, based on long-read metagenomic data from EP-1 and EP-2 samples, averaged by treatment. α-diversity of (A) ARGs and (B) MGEs measured using the Shannon index under different β-lactam treatments (Kruskal-Wallis *p < 0.05). (C) Normalized relative abundance of differentially abundant ARGs across treatments as measured with MaAsLin2 (qvalue < 0.05 |coefficient| > 1). Blue - antibiotic-free control, yellow - ampicillin, green - cefotaxime, orange - meropenem.

Overall, ampicillin supplementation exerted a limited effect on both ARG and MGE profiles. In contrast, exposure to cefotaxime and meropenem was associated with broader shifts, including a reduction in multiple *Enterobacteriaceae*-associated efflux pumps and MGEs. Specifically, several MGEs exhibited significant antibiotic-associated changes relative to no-AB controls: transposase-associated genes (*tnpA*-like) consistently decreased in abundance under all antibiotic treatments, while the insertion sequence *IS621* decreased under cefotaxime and meropenem exposure. In addition, the insertion element *ISBf10*, previously associated with *Bacteroides fragilis* (Kuwahara et al., 2004), showed reduced abundance under cefotaxime supplementation. In contrast, plasmid-associated replicons displayed antibiotic-specific increases, with *rep2* and *rep4* increasing under cefotaxime supplementation and *rep9, Col3M*, and *IncQ1* increasing under meropenem exposure. Notably, the class 1 integron integrase gene *intI1* decreased under meropenem supplementation (MaAsLin2 |coefficient| >1, qvalue < 0.05) (Supplementary table 1).

### Cefotaxime and meropenem exert different effects on the relative abundance and composition of enteric bacteria and plasmids

To acquire a more holistic understanding of how earlier and later β-lactams affect microbiome and resistome dynamics in gut-like communities, we assembled long-read sequencing data from both enrichment phases, generating large chromosome and plasmid contigs that can be more robustly annotated. Taxonomic and ARG profiling showed that many contigs carried genes conferring resistance to multiple antibiotic classes, with abundances varying by host taxa and resistance patterns (Fig. 5B), largely mirroring trends seen in the resistome and microbiome analyses detailed above. We identified several ARG-harboring contigs whose abundance was significantly different between the no-AB and the β-lactam-EP-2 samples. Collectively, the relative abundance of most assembled chromosomes and plasmids was lower in the cefotaxime- and meropenem-supplemented enrichments than in the no-AB and ampicillin-supplemented enrichments, and this was especially pronounced in gammaproteobacterial plasmids. Among the ESKAPE-E (*Enterococcus* spp., *Staphylococcus aureus, Klebsiella pneumoniae, Acinetobacter baumannii, Pseudomonas aeruginosa, Enterobacter* spp., and *Escherichia coli*) group, the relative abundance of *Enterococcus* was significantly higher in the meropenem-supplemented enrichment than in the no-AB, ampicillin- or cefotaxime-supplemented enrichments, whereas the relative abundance of *Escherichia* and other *Enterobacteriaceae* were significantly lower in cefotaxime and meropenem treatments relative to both the no-AB and the ampicillin-supplemented enrichments (Fig. 5A) (MaAsLin2 qvalue < 0.05, |coefficient| > 1).

**Fig 5:**
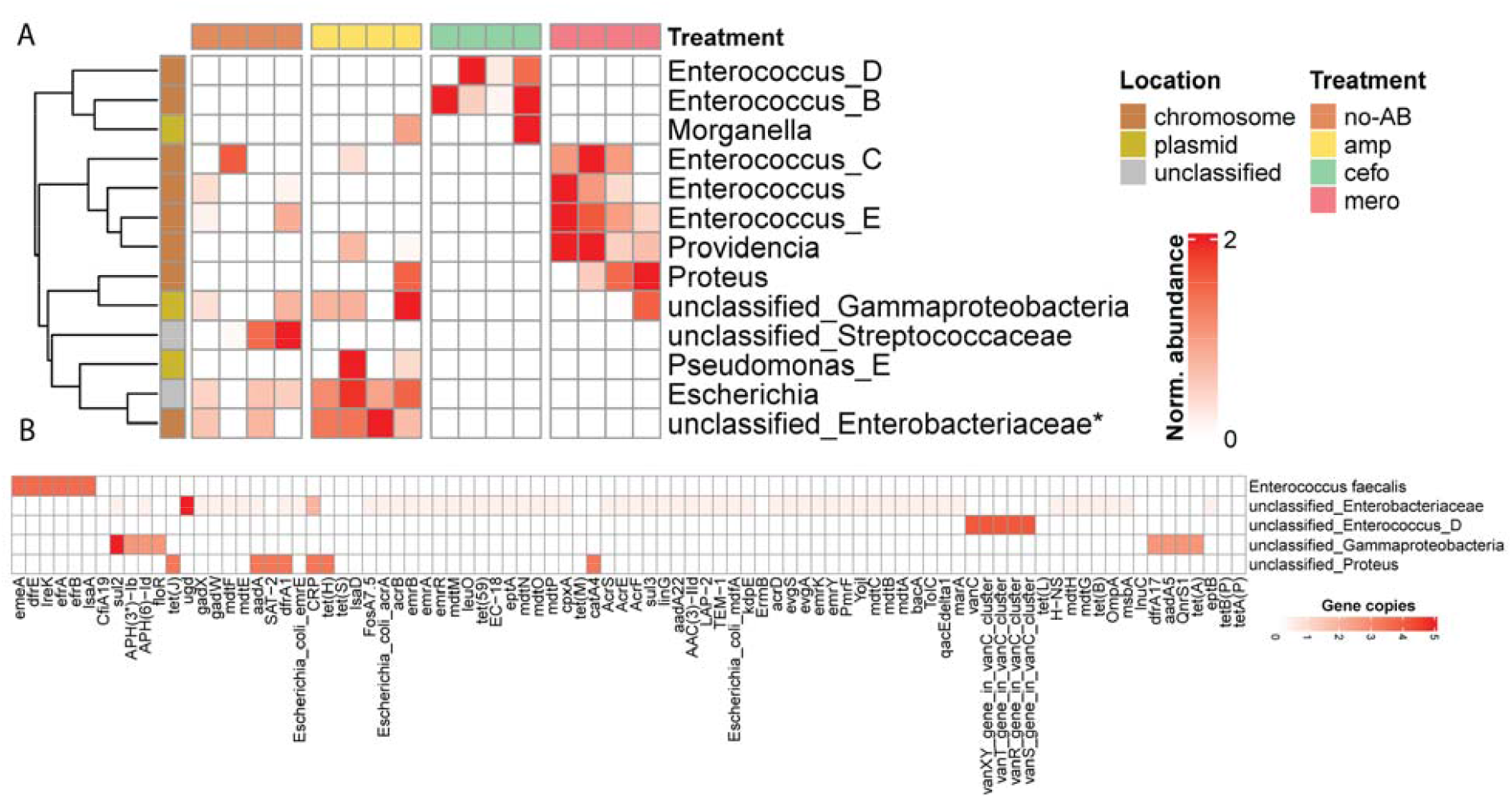
ARG-containing contigs that are differentially abundant between β-lactam and no-AB treatments in selective enrichments. (A) Normalized abundance of chromosome- and plasmid-associated contigs. (B) Characterization of ARGs in pathogen-associated taxa and plasmids (MaAsLin2; qvalue < 0.05 |coefficient| > 1). Letters adjacent to *Enterococcus* strains represent different assembled contigs.

## Discussion

Enrichment in BHI medium has been previously applied in several studies to establish stool-derived *in vitro* communities (SICs), following observations that the microbial composition of BHI-derived SICs largely mirrors that of their source stool (Aranda-Díaz et al., 2022; Goldman et al., 2025; Shi et al., 2025; Yousi et al., 2019a). BHI was recently implemented into an enrichment platform developed in our laboratory to identify gut-like bacteria in anthropogenically impacted produce (Davidovich et al., 2025). Here, we applied the BHI enrichment platform to chicken manure to examine the effects of early-versus later-introduced β-lactam antibiotics on gut-like microbiomes. The results of this study underline the fact that later β-lactams exert stronger selection on copiotrophic, gut-derived microbiomes than ampicillin. Nevertheless, the impact of different β-lactams is taxon-dependent, with distinct community members exhibiting differential responses to specific antibiotics, irrespective of their historical use.

Surprisingly, the microbiomes and resistomes of the ampicillin-supplemented enrichment strongly resembled the no-AB control, implying either that most of the copiotrophic gut-like taxa were resistant to ampicillin or that the proportion of extracellular β-lactamase producers was high enough to reduce ampicillin concentrations to below inhibitory levels (Wang et al., 2025). Nonetheless, ampicillin treatment resulted in lower relative abundances of *Lactococcus* and several *Clostridia*-associated genera compared with the no-antibiotic treatments, consistent with reports showing that while these β-lactamases are highly prevalent in *Enterobacteriaceae*, many Gram-positive species remain susceptible to older β-lactam antibiotics, potentially creating divergent impacts in mixed microbial communities during therapy (Ammor et al., 2007; Kakoullis et al., 2021a; Moore & Lacey, 2019; Toth et al., 2018).

Cefotaxime induced the largest shifts in microbial composition, notably affecting pathogenic-associate taxa such as *Escherichia–Shigella* and *Enterococcus*, while meropenem primarily reduced the abundance of several Bacillota genera, with *Escherichia–Shigella* being the only Pseudomonadota affected.

When evaluating bacterial proliferation under antibiotic selection, both intrinsic and acquired resistance must be considered (Belay et al., 2024; Mancuso et al., 2021), and the patterns observed here likely reflect the combined effects of intrinsic and acquired resistance across the taxa present in the manure-derived enriched samples, similar to previously reported studies (Udikovic-Kolic et al., 2014). For example, the increased relative abundances of *Enterococcus* strains in the cefotaxime- and meropenem-supplemented EP-2 samples relative to the ampicillin-supplemented and no-AB samples, can be explained by the fact that several of these strains are intrinsically resistant to cephalosporins and partially resistant to carbapenems (Iannetta et al., 2021; Kakoullis et al., 2021b), whereas *Escherichia–Shigella* mostly rely on acquired mechanisms for β-lactam resistance (Wang et al., 2025), which appeared to be absent in the enriched manure samples.

*Escherichia–Shigella* and other *Enterobacteriaceae* generally harbor a higher abundance of plasmids carrying mobile ARGs than other bacteria (Rozwandowicz et al., 2018), evident here in the observed diversity and relative abundance of ARGs and MGEs, which closely mirrored those seen in *Enterobacteriaceae*. The significant reduction in their relative abundance in cefotaxime- and meropenem-supplemented EP-2 enrichments compared to the no-AB and ampicillin treatments, is explained by the low abundance or absence of extended-spectrum β-lactamases and carbapenemases, respectively in the enriched plasmidome detected in this study, underlining the fact that these next-generation β-lactamases have not yet mobilized to poultry gut microbiomes.

Notably, most ARG-carrying plasmids were affiliated with Gammaproteobacteria and were depleted in the cefotaxime and meropenem treatments, consistent with the effects of these β-lactams on the *Escherichia–Shigella* genus. In addition, several chromosomal and plasmid fragments were linked to potentially pathogenic taxa and carried multiple ARGs, highlighting the potential risks associated with positive selection of these bacteria under antibiotic exposure.

The concomitant increase in relative abundance of specific enterococci species and the aminoglycoside resistance conferring gene *AAC(6*^*’*^*)-Iih* which has been previously reported to be harbored by enterococci (del Campo et al., 2005) in the cefotaxime-amended treatments, underlines the co-selection of intrinsic and acquired resistance mechanisms in the same bacteria as previously described (Simjee et al., 2024).

The increase in relative abundance of several taxa known to have intrinsic capacities in the cefotaxime and meropenem treatments might be explained by the depletion of antibiotic-sensitive competitors, highlighting how perturbations that reduce dominant taxa by antibiotics can lead to dramatic restructuring of gut communities (d’Humières et al., 2024; Martinson et al., 2019).

The study demonstrated the differential impact that antibiotics from the same group could have on microbiomes and resistomes in anthropogenic environments. While the application of antibiotics was previously reported to alter poultry gut microbiomes (Gupta et al., 2021; Kumar et al., 2018), these field results are often highly stochastic, difficult to replicate and therefore limited in their capacity to produce clear results. When coupled with field studies, the approach presented here where we specifically focused on an abridged copiotrophic community within the poultry gut microbiome can be applied to determine the impact of other antibiotics, coccidiostats or other xenobiotic compounds on poultry gut microbiomes. Furthermore, the methods can be used to evaluate the impact of various compounds on human gut microbiomes, and specifically dynamics of ESKAPE-E pathogens and various ARGs and MGEs.

While cefotaxime and meropenem are restricted to human use and are typically applied intravenously in hospitals, the detection of cefotaxime-resistant bacteria and associated ESBL-encoding genes in poultry manure, suggests that resistant bacteria and/or associated resistance-encoding plasmids have transferred to poultry via several plausible routes, including dust, human contact, feed, or wild avian sources (Erkorkmaz et al., 2025; Gonzalez-Martin et al., 2014; Hernandez et al., 2013). As demonstrated here, these transmission routes can affect the occurrence of pathogenic-associated strains even when the antimicrobial is applied to a non-target organism, potentially by selectively suppressing competing taxa within the microbial community (Letten et al., 2021).

When evaluating *Enterobacteriaceae* as a whole, and specifically *E. coli*, the fact that high levels of ampicillin resistance were observed vs. low levels of cefotaxime resistance and no resistance to meropenem, indicates that despite direct use, resistance to a given antibiotic develops as a function of the time, considering introduction of ampicillin, cefotaxime and meropenem in the early 1960’s, early 1980’s and mid-1990’s, respectively (Bush & Bradford, 2016).

Further experiments are needed to assess the impact of drug regimens, doses, and exposure times on microbiomes and resistomes for risk assessment, and to examine how these effects vary with sample source and native microbiome composition. pathogens that were susceptible two to three decades ago may now exhibit higher resistance levels, a trend that could continue with sustained antibiotic use and potentially accelerate progression toward a post-antibiotic era.

## Materials and Methods

### Manure sampling, enrichment, and gDNA extraction

Poultry manure was sampled from a battery hen poultry house located at the Volcani Institute, Israel, where chickens were fed with a standard feeding mix with 17% protein content. Manure samples were collected in sterile tubes from 4 different areas of the poultry house and transferred to the lab within 30 min, where 5 g from each manure sample was suspended in 30 ml 0.85% NaCl saline. Tubes were vortexed at max speed for 2 min and then transferred to a horizontal shaker at 500 RPM for 20 min to disengage bacteria from solid material into solution. Homogenates were then centrifuged for 10 min at 600 RPM at 4°C to settle particulate material and subsequently, 20 ml of supernatants were transferred to a new clean tube.

Phase one enrichments (EP-1) were prepared by inoculating 100 μl of the four above-described manure homogenates into 4.9 ml of Brain Heart Infusion (BHI) (Oxoid) in anoxic Hungate tubes (sparged with N_2_ to ensure O_2_ levels below 0.2 mg/L-measured with an O_2_ sensor (FT 55/12, YSI Inc., Ohio, USA)) using a sterile needle. BHI was chosen for enrichment due to previous reports showing that it allows the recovery of gut microbial communities (Lau et al., 2016; Yousi et al., 2019b). In tandem, 1 ml of manure homogenates was added to 750 μl of 50% glycerol for subsequent culture-based analyses. Enrichments were incubated for 20 hours at 180 rpm and 37° C, and glycerol stocks were prepared as described above. In addition, from each enrichment culture, 1□mL was transferred into a 1.5□mL Eppendorf tube and centrifuged at 5,000□×□g. The supernatant was discarded, and the bacterial pellets were stored at -80□°C for subsequent genomic DNA extraction.

Phase two enrichments (EP-2) were prepared by transferring 100 µl from each of the four EP-1 cultures into fresh anaerobic BHI medium, either without antibiotics (no-AB) or supplemented with ampicillin (8 ug/ml), cefotaxime (2 ug/ml), or meropenem (8 ug/ml), such that each of the four EP-1 replicates generated four corresponding EP-2 treatments (16 total). Antibiotic concentrations were selected based on EUCAST clinical breakpoints (version 11), using MIC breakpoints defined for Enterobacterales (EUCAST, 2021). EP-2 cultures were incubated for 20 hours at 180 rpm and 37° C, and processed similar to EP-1 enrichments.

For DNA extraction, EP-1 and EP-2 cultures were centrifuged for 1 minute at 5000 rpm, and supernatant was discarded. Pellets were extracted using the MagAttract High Molecular Weight (HMW) DNA extraction kit (Qiagen). DNA extraction from manure was performed using the DNeasy PowerMax Soil Kit (Qiagen), using 7 g of fresh manure, following the manufacturer’s protocol. The concentration of gDNA was determined using a Qubit fluorometer (Qubit 2.0, Invitrogen), and quality was assessed by measuring 260/280 and 230/260 ratios with the Nanodrop system 2000c Spectrophotometer (Thermo Fisher Scientific).

### Quantification, isolation, and identification of coliforms

*Escherichia-Shigella* counting was performed on Chromocult Coliform Agar plates (CCA; Merck, Darmstadt, Germany). EP-1 samples were serially diluted and then plated on CCA either without antibiotics or supplemented with ampicillin (8 µg/mL), cefotaxime (4 µg/mL), or meropenem (8 µg/mL). Concomitantly, EP-2 samples were plated on CCA without antibiotic selection. Cefotaxime-resistant *Escherichia-Shigella* were isolated from the cefotaxime-supplemented CCA plates for further analysis. Bacteria were initially characterized based on colony color on the CCA agar (blue or pink), and taxonomic affiliation was further validated by MALDI-TOF-MS (score > 2.0) using an Autoflex system following the direct colony method (Bruker Daltonics, Billerica, MA, USA). To avoid selection of identical clones for study, isolates were first genotyped using the Enterobacterial Repetitive Intergenic Consensus PCR (ERIC-PCR) method (Versalovic et al., 1991) prior to analysis. Selected isolates representing distinct genotypes were stored in glycerol stocks at -80 C°.

### 16S rRNA gene amplicon sequencing and data analysis

Native and enriched DNA samples were PCR amplified using the modified CS1-515F (GTGYCAGCMGCCGCGGTAA) and CS2-806R (GGACTAC-NVGGGTWTCTAAT) barcoded primers, which target the V4 region of the 16S rRNA gene, and sequencer-ready libraries were then generated as previously described [24]. The barcoded libraries were pooled and sequenced on a MiniSeq System (Illumina) at the Rush University DNA Services Facility, Chicago, Illinois, USA, using a 150-bp paired-end strategy with V3 chemistry. Paired-end reads were merged using PEAR (Zhang et al., 2014) and processed using QIIME2 (v2019.7) (Bolyen et al., 2019). The sequences underwent quality filtering employing the DADA2 algorithm (Callahan et al., 2016), which resolves amplicon sequence errors to generate amplicon sequence variants (ASVs). Taxonomic assignment was performed using the QIIME2 q2-feature-classifier with the SILVA 132 rRNA database (Quast et al., 2013). Samples with low sequencing depth, such as negative controls (< 1000 reads) were excluded from further analysis, as were ASVs corresponding to plastids or mitochondria. Raw count tables and the taxonomic assignments were then exported for further analyses and visualization in R.

### Long-read shotgun metagenome sequencing and analysis

Long-read metagenomic sequencing of EP-1 and EP-2 samples was performed on a Minion sequencer (Oxford Nanopore Technology (ONT), Oxford UK) using two Flongle flow cells (R9.4.1) in two independent 46 h runs. Basecalling was performed with the High Accuracy model in Guppy 6.0.7 (https://community.nanoporetech.com) with Qscore limit set to seven and min read length to 200 bp. *De novo* assembly and polishing (two polish cycles) were performed with metaFlye 2.6 (Kolmogorov et al., 2020) and Medaka 1.4.4 (https://github.com/nanoporetech/medaka), respectively. All bioinformatic analyses were performed using default parameters unless otherwise specified.

### Analysis of ARGs and MGEs in long-reads and assembled contigs

In order to annotate and quantify relative abundances of ARGs and MGEs, open reading frames (ORFs) were predicted from polished assemblies using Prodigal (v2.6.3) (Hyatt et al., 2010), and the resulting ORFs were merged into a combined set. Redundant ORFs were clustered with CD-HIT (v4.8.1) (Fu et al., 2012) at 90% identity to generate a non-redundant representative set. Annotation of predicted ORFs was performed against the SARG database for ARGs (Yin et al., 2023)and a custom MGE database (Pärnänen et al., 2018)using Diamond (v2.1.9) (Buchfink et al., 2021). Gene and read abundance in the metagenomes were estimated by mapping reads to the representative ORFs using Minimap2 (v2.17) (Li, 2018), followed by conversion of SAM files to BAM format, sorting, and indexing with Samtools (v1.9) (Li et al., 2009). Summary read counts for each ORF were obtained using samtools idxstats and further processed to generate final count tables. Polished contigs were annotated using Bakta (v1.9.2) (Schwengers et al., 2021) to generate standardized gene and functional annotations.

### Statistical analysis and visualization

Significant differences in alpha diversity (Shannon index) and beta diversity (Bray-Curtis, first and second PCoA axes) were assessed using linear mixed-effects models, with Treatment as a fixed effect and sample pairing as a random effect. Pairwise comparisons between treatments were performed with emmeans and Benjamini-Hochberg adjustment for multiple testing. All analyses were conducted in R using the lmerTest (v3.1.3) (Kuznetsova et al., 2017)and emmeans packages (v2.6.4) (https://github.com/rvlenth/emmeans). Differentially abundant taxa and genes were identified using the MaAsLin2 package (v1.2.0) (Mallick et al., 2021), and heatmaps were generated using the pheatmap package in R (v1.0.12) (Kolde, 2019).

Colony counting results were statistically analyzed using Dunnett’s test for multiple comparisons in JMP 18 and plotted with GraphPad Prism (v8.4.3).

