## Supplementary Data for "Extended-Spectrum and Last-Resort β-Lactams Exert Increasing Impacts on Chicken Manure-Derived Copiotrophic Microbiomes and Resistomes"

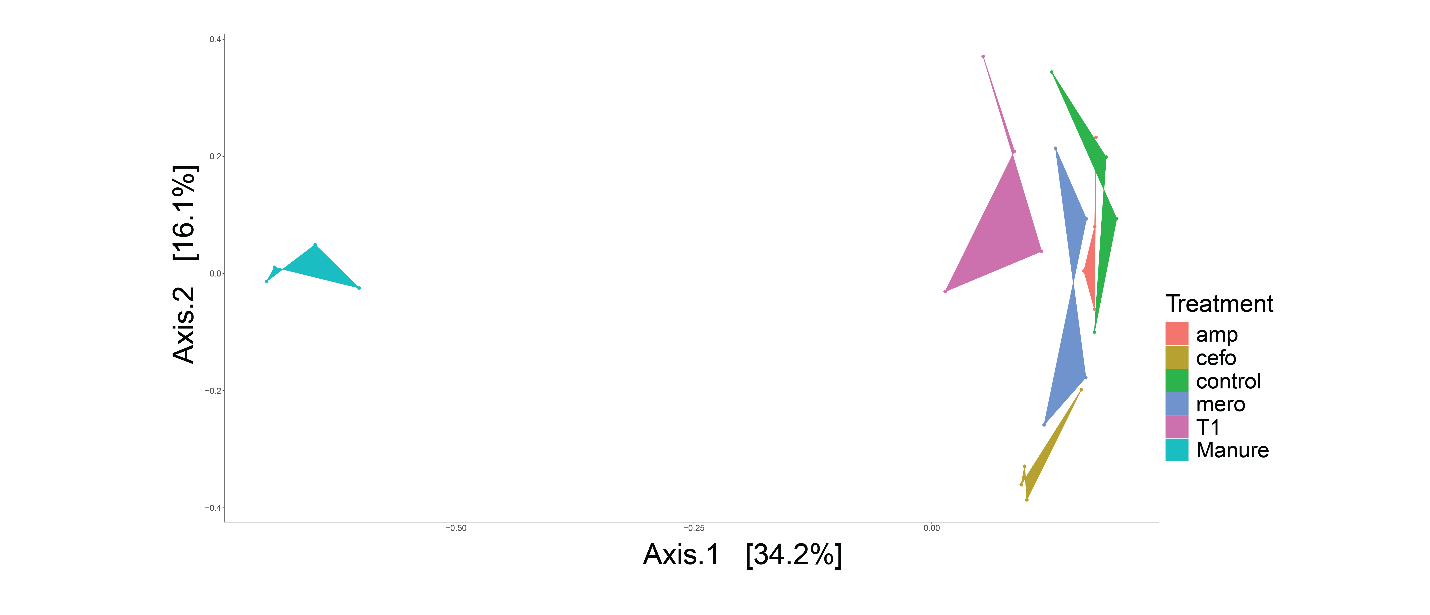

**Supplementary Fig. 1**. PCoA representation of β-diversity based on Bray-Curtis dissimilarity index of manure, non-selective (T1), and selective enrichments

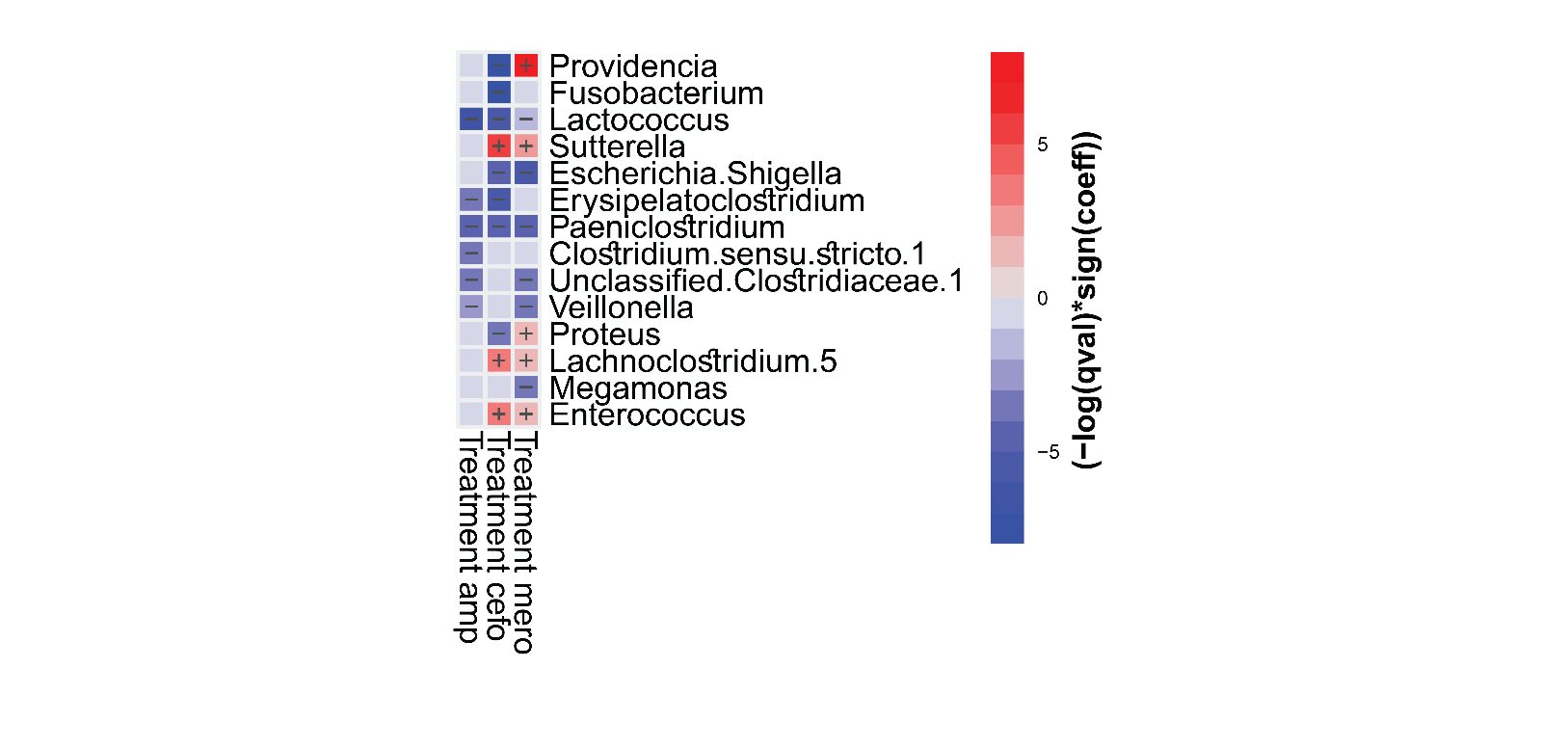

**Supplementary Fig. 2**. Differentially abundant genera between β-lactam-treated enrichments and no-AB as a reference. amp- Ampicillin; cefo- Cefotaxime; mero- Meropenem. MaAsLin2 qvalue < 0.05, |coefficient| > 1.

| feature | value | coef | stderr | qval |
| --- | --- | --- | --- | --- |
| tnpA1B1.ISLL6 | amp | -3.39549 | 0.391875 | 0.001972 |
| tnpA1B1.ISLL6 | cefo | -3.39549 | 0.391875 | 0.001972 |
| tnpA2B2.ISLL6 | amp | -3.88248 | 0.523579 | 0.003422 |
| tnpA2B2.ISLL6 | cefo | -3.88248 | 0.523579 | 0.003422 |
| tnpA1B1.ISLL6 | mero | -2.63679 | 0.391875 | 0.00581 |
| rep9 | cefo | -2.32474 | 0.418624 | 0.007069 |
| IS621 | mero | -4.14882 | 0.722877 | 0.011867 |
| tnpA2B2.ISLL6 | mero | -3.05837 | 0.523579 | 0.011867 |
| rep9 | amp | -2.06862 | 0.418624 | 0.012853 |
| tnpA3B3.ISLL6 | amp | -2.90308 | 0.566654 | 0.01927 |
| tnpA3B3.ISLL6 | cefo | -2.90308 | 0.566654 | 0.01927 |
| IS621 | cefo | -3.52565 | 0.722877 | 0.02281 |
| rep9 | mero | 1.845983 | 0.418624 | 0.02281 |
| rep22 | cefo | 3.0919 | 0.689443 | 0.03686 |
| tniA | cefo | -4.09894 | 0.953599 | 0.042277 |
| tniA | mero | -4.09894 | 0.953599 | 0.042277 |
| ISBf10 | cefo | -2.17288 | 0.516389 | 0.045468 |
| Col3M | mero | 2.996514 | 0.757644 | 0.047279 |
| IncQ1 | mero | 2.455413 | 0.60603 | 0.047279 |
| intI1 | mero | -2.32797 | 0.5789 | 0.047279 |
| ISSfl3 | cefo | -2.00004 | 0.512991 | 0.047279 |
| ISSfl3 | mero | -2.00004 | 0.512991 | 0.047279 |
| rep2 | cefo | 2.061533 | 0.51702 | 0.047279 |
| rep4 | cefo | 3.137116 | 0.795603 | 0.047279 |
| tnpA3B3.ISLL6 | mero | -2.33271 | 0.566654 | 0.047279 |
| tnpA4 | mero | -2.62464 | 0.652467 | 0.047279 |

**Supplementary Table 1**. MaAsLin2 analysis of MGEs that are differentially abundant between b-lactam enrichment and no-AB as a reference. Amp- Ampicillin; cefo- Cefotaxime; mero- Meropenem
